# Myofiber / pro-inflammatory macrophage interplay controls muscle damage in mdx mice

**DOI:** 10.1101/2020.12.28.424524

**Authors:** Marielle Saclier, Sabrina Ben Larbi, Eugénie Moulin, Rémi Mounier, Bénédicte Chazaud, Gaëtan Juban

**Affiliations:** Department of Biosciences, University of Milan, 20133 Milan, Italy; Institut NeuroMyoGène, Université Claude Bernard Lyon 1, CNRS UMR 5310, INSERM U1217, Université de Lyon, Lyon, France

**Keywords:** Duchenne muscular dystrophy, macrophage, fibrosis, myofiber branching

## Abstract

Duchenne Muscular Dystrophy is a genetic muscle disease characterized by chronic inflammation and fibrosis, which is mediated by a pro-fibrotic macrophage population expressing pro-inflammatory markers. The aim of this study was to characterize cellular events leading to the alteration of macrophage properties, and to modulate macrophage inflammatory status using the gaseous mediator H_2_S. We first analyzed the relationship between myofibers and macrophages in the mdx mouse model of Duchenne Muscular Dystrophy using coculture experiments. We showed that normal myofibers derived from mdx mice strongly skewed the polarization of resting macrophages towards a pro-inflammatory phenotype. Treatment of mdx mice with NaHS, an H_2_S donor, reduced the number of pro-inflammatory macrophages in skeletal muscle, which was associated with a decrease in the number of nuclei per fiber, a reduction of myofiber branching and a reduced fibrosis. These results identify an interplay between myofibers and macrophages where dystrophic myofibers contribute to the maintenance of a highly inflammatory environment that skews the macrophage status, which in turn favors myofiber damage, myofiber branching and fibrosis establishment. They also identify H_2_S donors as a potential therapeutic strategy to improve dystrophic muscle phenotype by modulating macrophage inflammatory status.

## Introduction

Adult skeletal muscle completely regenerates after an injury, thanks to the muscle stem cells (MuSCs) or satellite cells, that sustain adult myogenesis. This includes activation, proliferation, commitment into terminal myogenesis, differentiation and fusion to form new myofibers. The regeneration power of adult skeletal muscle is very high and damaged muscle fully recovers its functions within a few weeks. While MuSCs are absolutely indispensable for skeletal muscle regeneration, surrounding cells also play important roles in this process, including endothelial cells, fibro/adipogenic progenitors (FAPs) and immune cells (Bentzinger et al., 2013).

Particularly, macrophages are crucial for efficient muscle regeneration. As in other damaged tissues, the first macrophages to invade the injured area are pro-inflammatory damage-associated macrophages that mainly arise from circulating monocytes. Inflammatory damage-associated macrophages express pro-inflammatory effectors that stimulate MuSC proliferation (Saclier et al., 2013), while limiting FAP expansion (Juban et al., 2018; Lemos et al., 2015). Phagocytosis of debris participates to the resolution of inflammation, which is crucial to start tissue repair and regeneration (Arnold et al., 2007). Resolution of inflammation is characterized by a progressive switch of the inflammatory profile exhibited by macrophages, that become anti-inflammatory or recovery macrophages. This process is controlled by several molecular effectors, including the metabolic sensor AMPK (Mounier et al., 2013), the IGF1 pathway (Tonkin et al., 2016), the p38 regulator MKP-1 (Perdiguero et al., 2011), and the transcription factors C/EBPβ (Ruffell et al., 2009) and NFIX (Saclier et al., 2020). These anti-inflammatory macrophages exert a variety of effects on surrounding cells: they stimulate the last steps of myogenesis, including differentiation and fusion (Saclier et al., 2013), they activate FAPs to remodel the extracellular matrix (ECM) (Juban et al., 2018; Lemos et al., 2015), and favor angiogenesis, which occurs concomitantly to myogenesis (Latroche et al., 2017).

Post-acute injury skeletal muscle regeneration is a highly regulated process, during which the various stages described above must be well coordinated in space and time. The situation is drastically different in degenerating myopathies, such as Duchenne Muscular Dystrophy (DMD), during which multiple injuries occur asynchronously throughout the muscle. Skeletal muscle in degenerative myopathies fails to regenerate because of these permanent and asynchronous cycles of degeneration-regeneration due to mutations in the dystrophin-glycoprotein complex, which ensures myofiber integrity (Lapidos et al., 2004). In these disorders, recurrent myofiber damage leads to chronic inflammation and the eventual establishment of fibrosis. Indeed, it was shown that asynchronous injuries in normal muscle trigger a chronic inflammatory status in the muscle associated with a fibrotic signature, reminiscent of what is observed in degenerative myopathies (Dadgar et al., 2014).

Macrophages are associated with fibrosis in mdx muscle, the mouse model for DMD (Vidal et al., 2008). Inhibition of the pro-inflammatory NF-κB pathway is associated with a lower macrophage number and a better phenotype of the muscle (Acharyya et al., 2007). Similarly, preventing the entry of circulating monocytes into the muscle temporarily improves muscle histology and function (Liang et al., 2018; Mojumdar et al., 2014; Wehling et al., 2001). However this is detrimental on a longer term (Zhao et al., 2016), showing the pivotal role of macrophages during muscle regeneration, and the necessity to target the right subset to prevent disease progression. Recently, we showed in an experimentally fibrotic mdx mouse model, that fibrosis is associated with pro-inflammatory macrophages (Juban et al., 2018). Functionally, these macrophages stimulate matrix production by fibroblastic cells through the abnormal secretion of latent TGFβ, due to an increased expression of the gene encoding a Latent TGF-beta binding protein (LTBP), that is involved in latent TGFβ export into the ECM. Interestingly, pharmacological skewing of these macrophages towards an antiinflammatory phenotype using an AMPK activator improves muscle phenotype and function (Juban et al., 2018).

The standard treatment for DMD patients consists of Glucocorticoids (GCs), which have been shown to delay disease progression and ambulation loss (McDonald et al., 2018). However, chronic daily exposure to GCs is associated with adverse effects including obesity, metabolic changes and muscle atrophy (Schakman et al., 2013; Wood et al., 2015). Therefore, alternative therapeutic strategies are required. Hydrogen Sulfide (H_2_S) is an endogenously synthetized small gaseous signaling molecule that is freely permeable to cell membrane (Wang, 2014). H_2_S is involved in various biological processes, including inflammation. Alteration of H_2_S levels have been associated with several diseases in human or animal models, including sepsis, diabetes or rheumatoid arthritis (Fagone et al., 2018). H_2_S has emerged as a promoter of the resolution of inflammation through the regulation of macrophage inflammatory state (Sun et al., 2020; Wallace et al., 2012). Interestingly, H_2_S-releasing molecules, like NaHS, have been shown to reduce skeletal muscle atrophy in a mouse model of diabetes (Lu et al., 2020) and to decrease muscle fibrosis after contusion-induced injury (Zhao et al., 2020). Moreover, H_2_S-releasing molecules have shown promising results in cardiovascular disorders and arthritis with reduced adverse effects (Wallace et al., 2020; Wallace et al., 2018).

In the present study, we investigated the interaction between myofibers and macrophages in the context of degenerative myopathies. As the macrophage phenotype is modulates by their close environment, we tested the hypothesis that dystrophic muscle fibers may themselves alter macrophage status, using *ex vivo* coculture experiments. Then, we analyzed the possibility to modulate the macrophage phenotype *in vivo* using NaHS, an H_2_S donor, to ameliorate dystrophic muscle phenotype.

## Results

### DMD-derived myofibers trigger pro-inflammatory profile of macrophages

The role of myofibers on macrophage polarization was investigated by coculturing wildtype bone marrow derived macrophages (BMDMs) with single fibers isolated from wild-type and dystrophic muscle (Fig.1A). Wild-type myofibers increased the expression of 2 out of 4 pro-inflammatory markers analyzed by resting macrophages (Fig.1B-C, +23% for TNFα and +37% for COX2 as compared with control). However, in the presence of mdx myofibers, the number of BMDMs expressing pro-inflammatory markers was strongly increased: it increased the expression of 3 out of 4 pro-inflammatory markers: TNFα, CCR2 and COX2 (Fig.1B-C, 45, 30 and 58% increase as compared with control, respectively). No modification of the expression of the anti-inflammatory markers CD301 and ARG1 by BMDMs was observed when cocultured with wild-type or mdx myofibers (Fig.1B-C). These results show that mdx myofibers skew macrophages towards a pro-inflammatory phenotype. Thus, not only the cycles of injury/regeneration, but also the dystrophic myofibers themselves sustain the inflammation in mdx muscles, highlighting the relevance of modulating inflammation as a therapeutic strategy.

**Figure 1.**
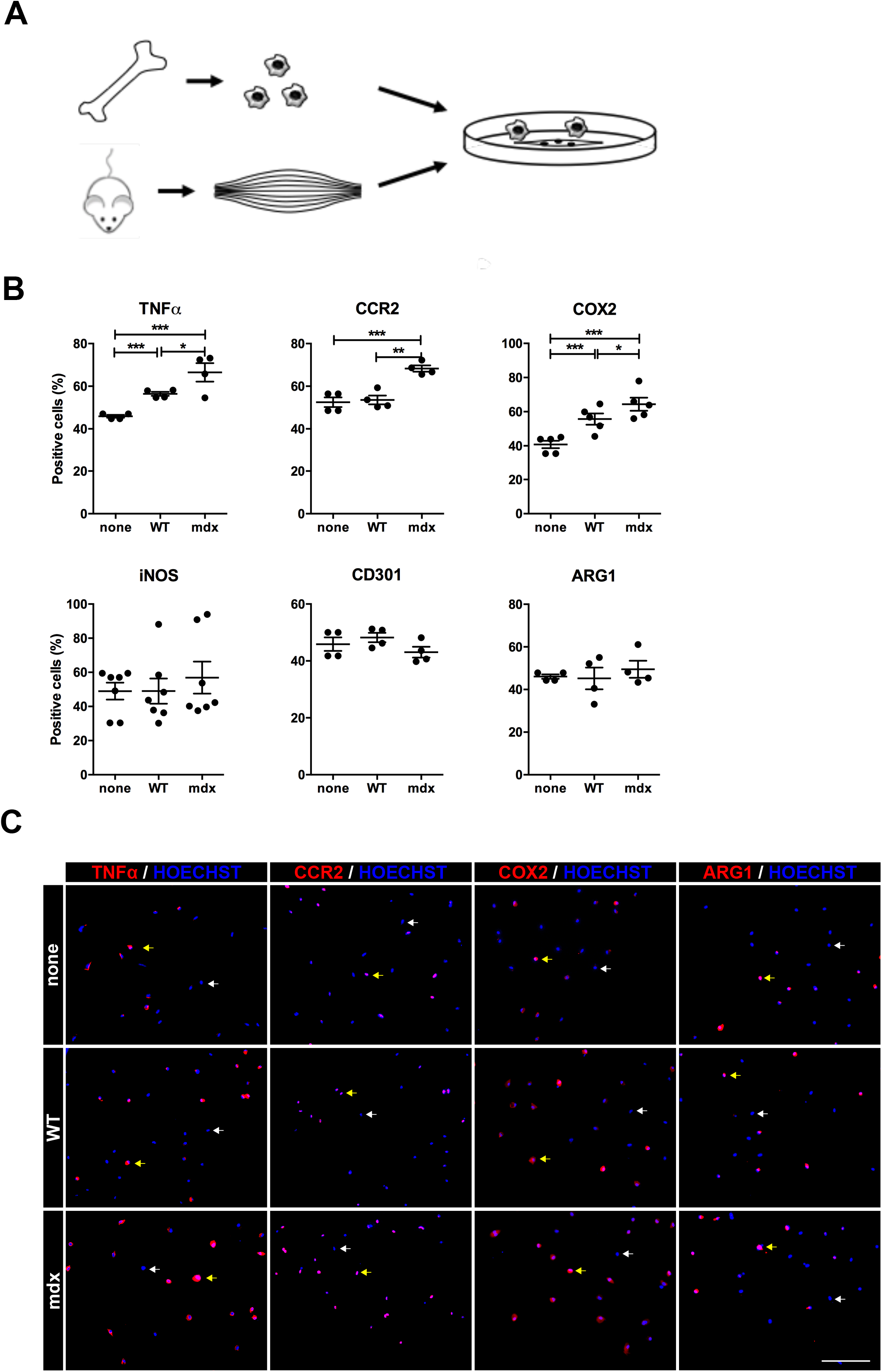
Dystrophic myofibers trigger pro-inflammatory macrophages. **(A)** Diagram depicting the cocultures of BMDMs with isolated single myofibers. **(B-C)** BMDMs that were cultured alone (none) or with wild-type (WT) or mdx myofibers for 3 days were immunolabelled for pro- (TNFα CCR2, COX2 and iNOS) or anti- (CD301, ARG1) inflammatory markers. **(B)** Percentage of macrophages positive for the specified marker. **(C)** Representative images of TNFα, iNOS, COX2 and ARG1 immunolabelings. Scale bar, 50 μm. Arrows indicate macrophages negative (white arrow) or positive (yellow arrow) for the specified marker. Results are means ± sem of 4-7 experiments. *p<0.05; **p<0.01; ***p<0.001.

### NaHS treatment reduces pro-inflammatory macrophages in dystrophic muscle

In an attempt to modulate the macrophage inflammatory status, we treated 3-month-old mdx mice with NaHS, an H_2_S donor. Because of the short half-life of H_2_S *in vivo* (Calvert et al., 2010), NaHS was delivered intraperitoneally daily for 3 weeks (Fig.2A). In the *Tibialis Anterior* (TA) muscle, NaHS treatment induced an important decrease in the total number of macrophages (Fig.2B, −42%). More precisely, in NaHS treated mice, a strong decrease in the number of macrophages expressing pro-inflammatory markers was observed (Fig.2C-D, −35 and −31% for TNFα and iNOS, respectively) together with a decrease of the fibrosis-associated ARG1 marker (Fig.2C-D, −35%), and a trend towards a slight increase of the anti-inflammatory marker CD206 (Fig.2C, +17%). The expression of CCR2 and CD301 was not affected (Fig.2C). These results show that NaHS treatment specifically reduces the number of pro-inflammatory macrophages in mdx mice muscle.

**Figure 2.**
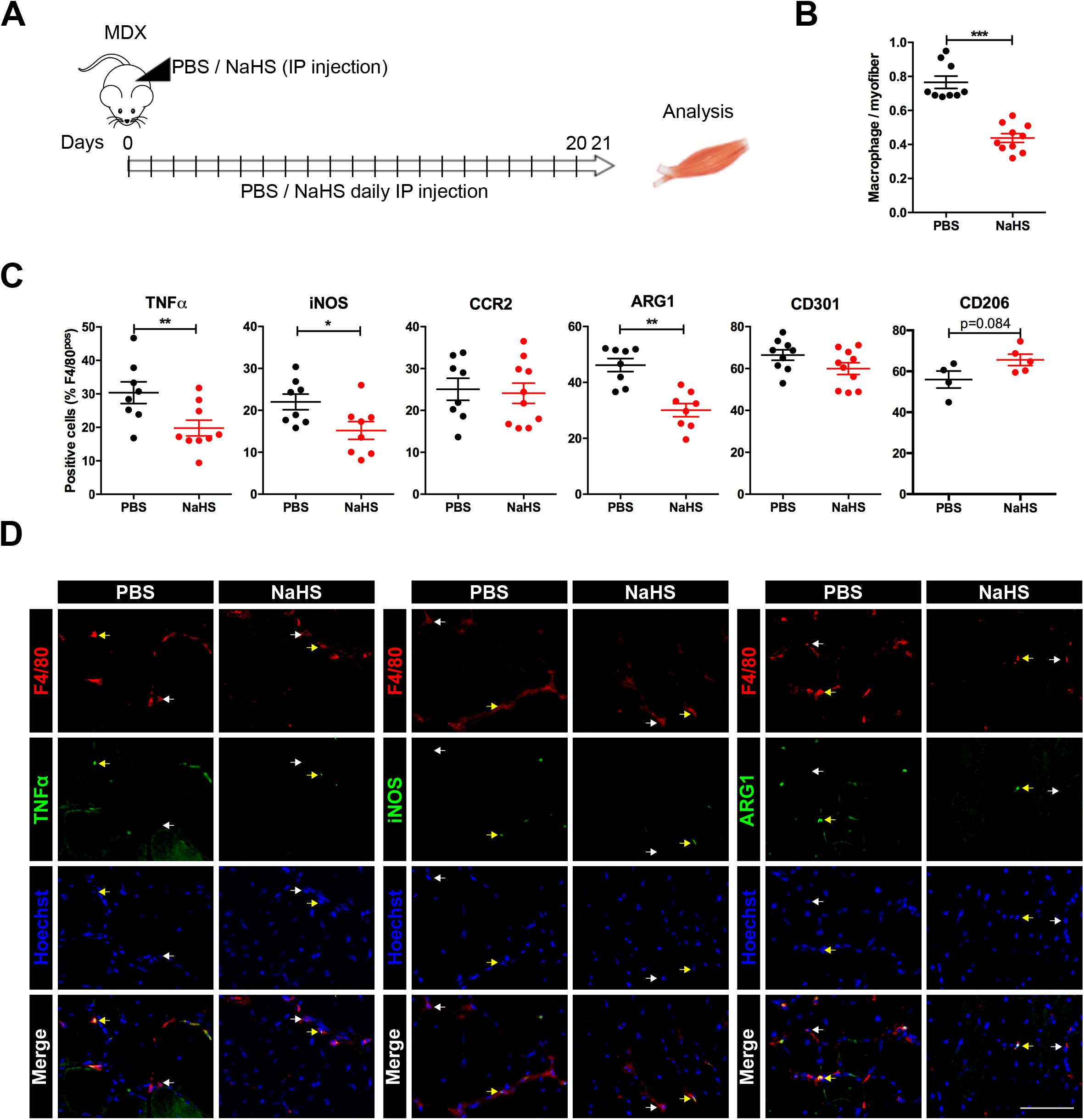
NaHS treatment reduces pro-inflammatory macrophages in mdx muscle. **(A)** Mdx mice were treated daily with PBS or NaHS for 3 weeks and *TA* muscles were harvested. **(B)** Muscle sections were immunostained for F4/80 and Laminin and the number of macrophages per myofiber was determined. **(C-D)** Muscle sections were immunolabeled for F4/80 and pro- (TNFα, iNOS, CCR2) or anti- (CD301, CD206, ARG1) inflammatory markers. **(C)** Percentage of macrophages positive for the specified marker. **(D)** Representative images of F4/80 and TNFα (left panel), iNOS (middle panel) and ARG1 (right panel) immunolabelings. Arrows indicate macrophages negative (white arrow) or positive (yellow arrow) for the specified marker. Scale bar, 50 μm. Results are means ± sem of 4-10 experiments. *p<0.05; **p<0.01; ***p<0.001.

### Pharmacological decrease of macrophage pro-inflammatory phenotype by NaHS reduces myofiber branching in mdx mice

To study the impact of pro-inflammatory macrophage reduction in dystrophic muscle, several myogenic parameters were analyzed after NaHS treatment. Immunostainings performed on TA transversal sections from treated mdx muscles did not show a modification of the number of satellite cells per fiber as determined by Pax7 immunostaining (Fig.3A), or of the proportion of newly formed myofibers that expressed eMHC+ (Fig.3B). However, NaHS treatment led to a decreased number of nuclei per myofiber (Fig.3C-D, 20% decrease), indicating a reduction in cell fusion. Muscle regeneration after an injury is associated with myofiber branching (Pichavant and Pavlath, 2014), which is characterized by myofibers containing two or more cytoplasmically continuous strands. Mdx mice harbour an increased proportion of branched myofibers as compared with wild-type (Bockhold et al., 1998), likely due to the permanent cycles of injury/regeneration, which is correlated with a decrease in muscle force production (Chan et al., 2007). Interestingly, isolated myofibers from NaHS-treated mdx mice were characterized by a decrease in myofiber branching (Fig.3E-F, 18% decrease). Altogether, these results show that dampening the pro-inflammatory status of macrophages is associated with a reduction of myofiber branching and of the number of myonuclei/myofiber, indicative of a reduction of muscle damage.

**Figure 3.**
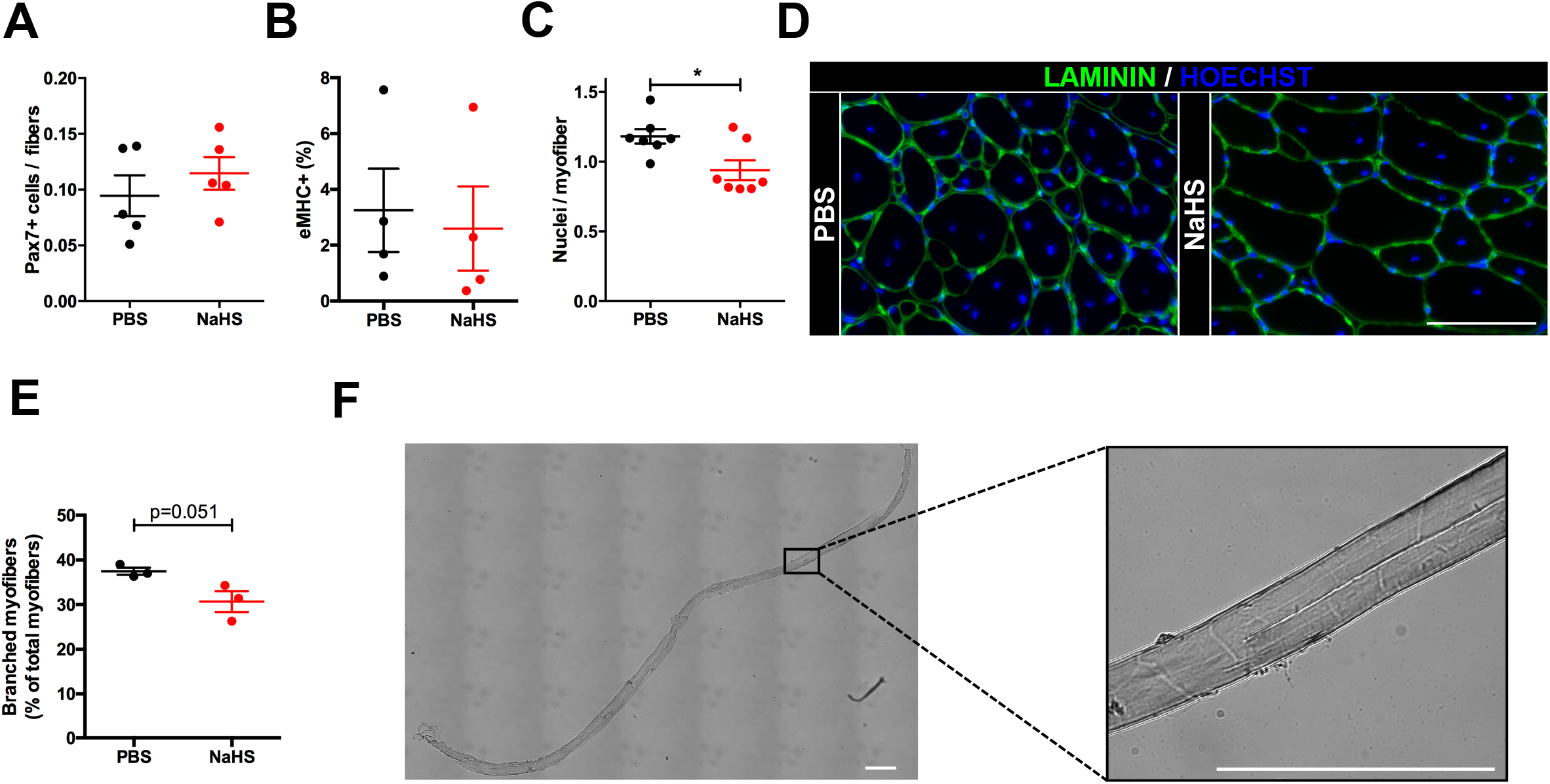
NaHS treatment reduces muscle damage in mdx mice. Mdx mice were treated daily with PBS or NaHS for 3 weeks (as described in Fig.2A), TA muscles were harvested and immunostainings were performed. **(A)** Number of satellite cells (as Pax7^pos^ cells) normalized per the number of myofibers. **(B)** Newly formed myofibers (as eMHC^pos^ myofibers), represented in percent of the total number of myofibers. **(C-D)** Muscle sections were immunolabeled for Laminin and Hoechst to determine the number of nuclei per myofiber. **(C)** Number of nuclei per myofiber. **(D)** Representative images of Laminin and Hoechst immunolabelings. Scale bar, 50 μm. **(E-F)** Single myofibers were isolated to determine the proportion of branched myofibers. **(E)** Percentage of branched myofibers. **(F)** Representative image of a branched myofiber. Scale bar, 100 μm. Results are means ± sem of 3-7 experiments. *p<0.05.

### NaHS treatment improves dystrophic muscle

Next, we evaluated the consequence of dampening the macrophage pro-inflammatory status by NaHS treatment on dystrophic muscle features. Hematoxylin-eosin (HE) staining of TA transversal sections showed a two fold decrease in the damaged areas in NaHS-treated mice (Fig.4A-B, −50%). This was associated with a decrease in muscle fibrosis, as shown by the reduction in the areas expressing Collagen1 (Coll1) (Fig.4C-D, −10.4%). Finally, the mean cross-sectional area (CSA) of myofibers, which is correlated with muscle force production, was increased (Fig.4E-F, +18%). Precisely, a decreased proportion of the smallest fibers (Fig.4E-F, −28% of fibers with CSA <500 μm^2^) and a concomitant increased proportion of the biggest fibers (Fig.2F-G, +17 and +23% of fibers with CSA between 3000-3500 and 3500-4000 μm^2^, respectively) were observed in NaHS-treated mdx mice. Taken together, these results show that the reduction of pro-inflammatory macrophages by NaHS treatment is associated with an amelioration of the dystrophic muscle phenotype.

**Figure 4.**
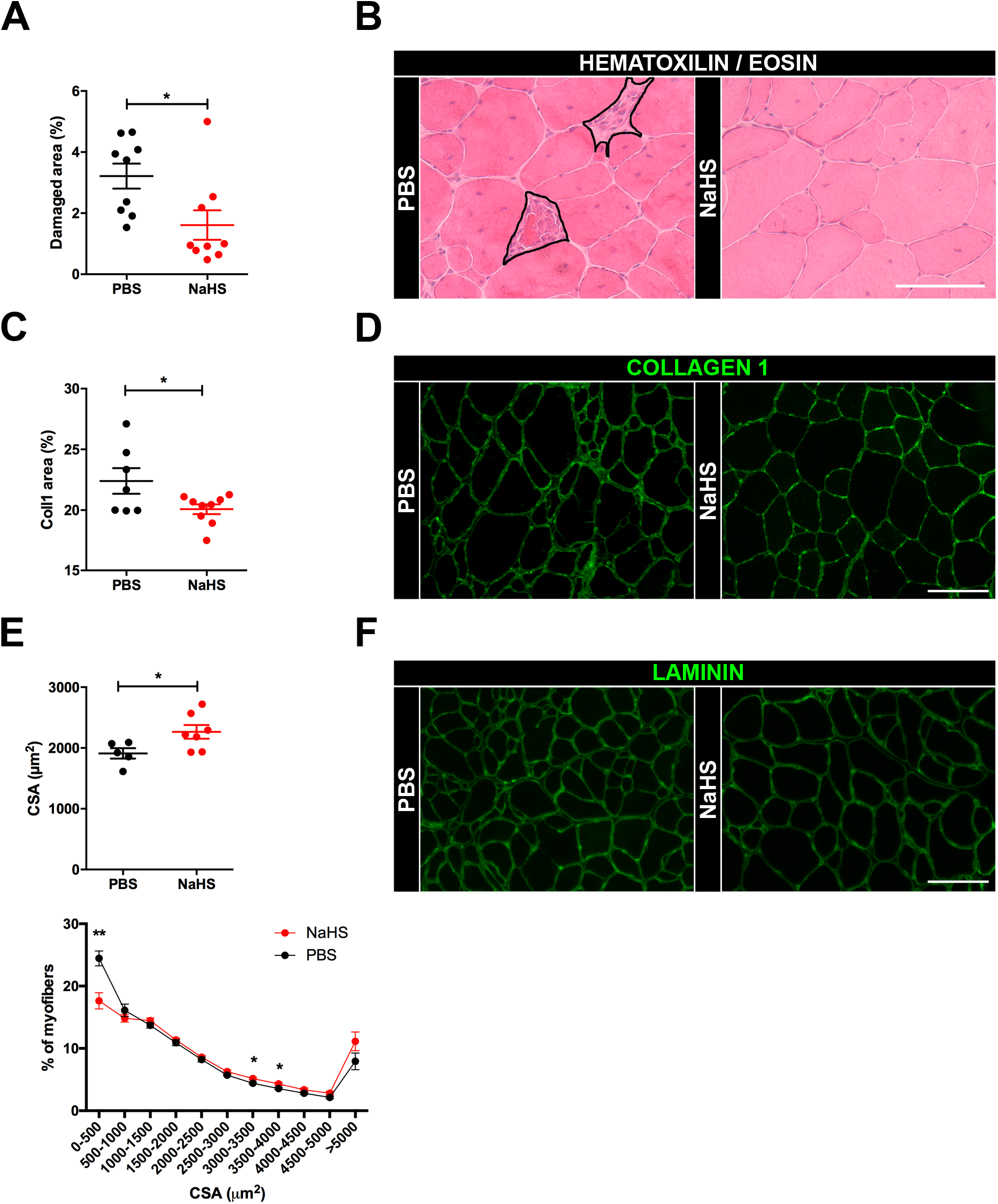
NaHS treatment ameliorates dystrophic muscle phenotype. Mdx mice were treated daily with PBS or NaHS for 3 weeks (as described in Fig.2A) and TA muscles were harvested. **(A-B)** Muscle sections were stained with HE and the damaged area were determined. **(A)** Percentage of damaged area per muscle section. **(B)** Representative images of HE staining showing damaged areas circled in black. Scale bar, 50 μm. **(C-D)** Muscle sections were immunostained for Coll1. **(C)** Percentage of Coll1 area. **(D)** Representative images of Coll1 immunostaining. Scale bar, 50 μm. **(E-F)** Myofiber Cross-Section Area (CSA) was determined on whole muscle section after Laminin immunolabeling. **(E)** Mean CSA (top graph) and repartition of myofibers by size (bottom graph). **(F)** Representative images of Laminin immunolabeling. Scale bar, 50 μm. Results are means ± sem of 5-9 experiments. *p<0.05; **p<0.01.

## Discussion

In the present study, we analyzed the interaction between myofibers and macrophages in the context of DMD. We showed that the dystrophic myofibers themselves skew resting macrophages towards a pro-inflammatory phenotype. As alterations in the secretome of myotubes generated by myoblasts isolated from mdx mice (Duguez et al., 2013) or DMD patients (Lecompte et al., 2017) have already been described, this suggests that in the context of degenerative myopathies, not only the degenerating/regenerating events lead to chronic inflammation, but the myofibers themselves contribute to the maintenance of the pro-inflammatory environment that alters macrophage properties and favors the generation of a pro-fibrotic population. Indeed, we previously showed that pro-inflammatory macrophages promote fibrosis in DMD muscle (Juban et al., 2018). Moreover, modulating the inflammatory status of this population towards an anti-inflammatory phenotype improves DMD muscle function (Juban et al., 2018). Therefore, identification of specific anti-inflammatory strategies appears as a relevant therapeutic approach to alleviate muscle damage.

H_2_S is a gasotransmitter that exhibit anti-inflammatory properties, in part through the regulation of macrophage inflammatory status (Sun et al., 2020; Wallace et al., 2012). H_2_S donors have been widely used *in vivo* using animal models to treat various pathological conditions (Fagone et al., 2018). This includes reduction by the H_2_S donor NaHS of skeletal muscle atrophy in a mouse model of diabetes (Lu et al., 2020), or the decrease of muscle fibrosis after contusion-induced injury (Zhao et al., 2020). Interestingly, various therapeutics like Zofenopril have been shown to exert their effect through the indirect release of H_2_S (Fagone et al., 2018; Wallace et al., 2018). Recently, clinical studies have shown the safety and efficiency of H_2_S-releasing molecules in human. Patients with congestive heart failure exhibit a deficit of H_2_S in the blood. Treatment with the H_2_S donor SG1002 attenuated the level of brain natriuretic peptide, a stress marker of cardiac function, thus reducing the severity of and potentially preventing heart failure (Polhemus et al., 2015). Similarly, ATB-346, an H_2_S-releasing derivative of the nonsteroidal anti-inflammatory drug Naproxen, reduced the symptoms severity in osteoarthritis patients with reduced adverse effects as compared to Naproxen alone (Wallace et al., 2018; Wallace et al., 2020).

At the molecular level, the H_2_S donor NaHS was shown to favor microglia inflammatory shift towards an anti-inflammatory phenotype (Du et al., 2014; Zhou et al., 2014) through the activation of AMPK (Zhou et al., 2014). Moreover, its effect on the down-regulation of pro-inflammatory markers expression in BMDMs following LPS stimulation requires the resolving Annexin A1 (Brancaleone et al., 2014). Interestingly, our previous work identified AMPK as a crucial activator of macrophage inflammatory shift during normal skeletal muscle regeneration (Mounier et al., 2013). Additionally, we showed that Annexin A1 activates AMPK in macrophages in this context (McArthur et al., 2020).

Thus, in an attempt to counteract the exacerbated macrophage pro-inflammatory phenotype induced by dystrophic myofibers, we treated mdx mice with NaHS. We showed that NaHS-treated mdx mice exhibited a decrease in the total number of macrophages. More specifically, we observed a reduction in the number of macrophages expressing pro-inflammatory markers as well as the ARG1 fibrotic marker. This was associated with a global muscle phenotype amelioration characterized by a reduced myofiber necrosis and reduced fibrosis, together with an increased myofiber size. This is in accordance with the beneficial effect observed on dystrophic muscle function by preventing the entry of circulating monocytes at early time points (Liang et al., 2018; Mojumdar et al., 2014; Wehling et al., 2001), or by skewing macrophage towards an anti-inflammatory phenotype (Juban et al., 2018). These results confirm that the modulation of macrophage inflammatory status is a relevant therapeutic approach to relieve inflammation and muscle damage in degenerative myopathies.

The amelioration of the muscle phenotype by NaHS treatment was characterized by a decrease in the number of nuclei per myofiber together with a reduction of myofiber branching. Branched myofibers are barely observed in normal skeletal muscle but they appear after regeneration following acute injury (Pichavant and Pavlath, 2014). They are observed in DMD patients (Bell and Conen, 1968) and in DMD fibers generated *in vitro* (Tanoury et al., 2020), as well as in the mdx mouse model (Head et al., 1992; Lefaucheur et al., 1995). Branched myofibers are characterized by an impairment of calcium signaling and Excitation-Contraction coupling, causing myofiber function defects and higher fragility (Head, 2010; Lovering et al., 2009). While the mechanisms resulting in myofiber branching are still unclear (defect in final fusion between forming myofibers, or normal process that is exacerbated in dystrophic muscle due to continuous regeneration processes), a recent study showed in the mdx EDL muscle that the increased number of myonuclei per myofiber is concomittant to myofiber branching (Massopust et al., 2020). Accordingly, the concomitant reduction of the number of nuclei per myofiber and of branching observed in NaSH-treated mice suggests an improvement in muscle homeostasis.

To conclude, the present study identified a novel property of dystrophic myofibers in the DMD mouse model mdx, associated with a regulatory loop between myofibers and macrophages. Dystrophic myofibers skew macrophages towards a pro-inflammatory phenotype that in turn contributes to myofiber damage, consequently increasing myofiber branching and fragility, thus favoring muscle damage and fibrosis. Finally, this work identified H_2_S-releasing molecules as potential therapeutic strategies to dampen inflammation in DMD and improve dystrophic muscle phenotype.

## Materials and methods

### Mice experiments

DMD^mdx4Cv^ (Chapman et al., 1989) mice on C57BL/6J background were bred and used according to French legislations. Experiments were conducted on males at 10-12 weeks of age. NaHS (1 mg/kg, Sigma-Aldrich) or PBS was injected daily intraperitoneally in mdx mice for 3 weeks and muscles were harvested the day after the last injection.

### Histology and immunofluorescence analyses in mouse

Fascia of TA muscles was removed, then muscles were frozen in nitrogen-chilled isopentane and kept at −80°C until use. 8 μm-thick cryosections were prepared for HE staining, and immunolabelings. For immunolabelings, cryosections were permeabilized 10 min in 0.5 % Triton X-100 and saturated in 2% BSA for 1 h at room temperature. For identification of regenerating myofibers, saturated cryosections were co-labeled with primary antibodies directed against embryonic myosin (eMHC, sc-53091, Santa Cruz) and Laminin (L9393, Sigma Aldrich). For macrophage double immunolabelings, cryosections were labeled with antibodies (from Abcam unless indicated) against F4/80 (ab6640, ab74383) overnight at 4°C and labeling using the second antibody was performed for 2 h at 37°C. Antibodies were directed against Arginase 1 (sc-18355, Santa Cruz), CCR2 (ab32144), CD206 (sc-58987, Santa Cruz), CD301 (ab59167), Collagen I (ab292, ab34710 or Biotech 131001), Cox2 (ab2372), iNOS (ab15323), Laminin (L9393, Sigma Aldrich, or ab78287) and TNFα (ab34839). Secondary antibodies were coupled to FITC, Cy3 or Cy5 (Jackson Immunoresearch Inc). Muscle stem cells were labeled as previously described (Théret et al., 2017) using an antibody directed against Pax7 (AB_528428, DSHB). HE-stained muscle sections were recorded with a Nikon E800 microscope at 20X magnification connected to a QIMAGING camera. Damaged areas were measured as previously described (Juban et al., 2018). Fluorescent immunolabelings were recorded with a DMI 6000 Leica microscope connected to a Coolsnap camera at 20X magnification. For each condition of each experiment, at least 8-10 fields chosen randomly were counted. The number of labeled macrophages was calculated using the Image J software and was expressed as a percentage of total macrophages. Areas of Collagen1 were calculated with ImageJ software as previously described (Juban et al., 2018). Cross Section Area was determined on whole muscle sections labeled by laminin antibody using Open-CSAM program as previously described (Desgeorges et al., 2019).

### BMDMs

Macrophages were derived from murine bone marrow precursors as previously described (Mounier et al., 2013). Briefly, total bone marrow was obtained by flushing femur and tibiae from wild-type C57BL/6J mice with Dubecco’s Modified Eagle’s Medium (DMEM). Cells were cultured in DMEM containing 20% Fetal Bovine Serum (FBS), 30% of L929 cell line-derived conditioned medium (enriched in CSF-1), 2.5 μg/mL of fungizone and 100 U/mL of penicillin for 6-7 days.

### Cocultures of macrophages with isolated single myofibers

Single myofibers of *extensor digitorum longus* (EDL) muscle were isolated from wild-type and mdx mice as previously described (Le Grand et al., 2012) and were cocultured as 15-20 myofibers/well in 96 wells containing 14000 cells/cm^2^ of wild-type bone-marrow derived macrophages (prepared as described above) for 3 days. Macrophages were immunolabelled as described above for the immunofluorescence on muscle sections. For each condition of each experiment, about 10-12 fields chosen randomly were recorded with a DMI 6000 Leica microscope connected to a Coolsnap camera (Photometrics) at 20X magnification.

### Isolation of single myofibers from TA muscle

Fascia of TA muscles was removed and muscles were fixed for 45 min in 4% formaldehyde. After 2 quick washes and a 30 min wash in PBS, muscles were transferred into PBS containing 0.5% Triton X-100. 30 to 40 single myofibers were gently isolated by manual dissociation using tweezers and were mounted on slides. Slides were observed on an Axio Observer.Z1 (Zeiss) connected to a CoolSNAP HQ2 CCD Camera (Photometrics) and the branched myofibers were numerated.

### Statistical Analyses

All experiments were performed using at least three different cultures or animals in independent experiments. The Student’s t test was used for statistical analyses. P < 0.05 was considered significant.

## Acknowledgements

This work was funded by the Framework Programme FP7 Endostem (under grant agreement 241440), Association Française contre les myopathies (MyoNeurAlp Alliance) and Fondation pour la Recherche Médicale (Equipe FRM DEQ20140329495).

## Author contribution

Conceptualization: GJ, BC, RM; Methodology: MS, SBL, GJ, EM, BC; Analysis: SBL, GJ, MS, BC; Writing original draft: GJ, MS; Writing review & editing: GJ, MS, BC, RM; Supervision: GJ, BC; Funding Acquisition: BC.

